# Reconstruction of phylogenetic history to resolve the subspecies anomaly of Pantherine cats

**DOI:** 10.1101/082891

**Authors:** Ranajit Das, Priyanka Upadhyai

## Abstract

All charismatic big cats including tiger (*Panthera tigris*), lion (*Panthera leo*), leopard (*Panthera pardus*), snow leopard (*Panthera uncial*), and jaguar (*Panthera onca*) are grouped into the subfamily Pantherinae. Several mitogenomic approaches have been employed to reconstruct the phylogenetic history of the Pantherine cats but the phylogeny has remained largely unresolved till date. One of the major reasons for the difficulty in resolving the phylogenetic tree of Pantherine cats is the small sample size. While previous studies included only 5‐10 samples, we have used 43 publically available taxa to reconstruct Pantherine phylogenetic history. Complete mtDNA sequences were used from all individuals excluding the control region (15,489bp). A Bayesian MCMC approach was employed to investigate the divergence times among different Pantherine clades. Both maximum likelihood and Bayesian phylogeny generated a dendrogram: *Neofelis nebulosa* (*Panthera tigris* (*Panthera onca* (*Panthera uncia* (*Panthera leo*, *Panthera pardus*)))), grouping lions with leopards and placing snow leopards as an outgroup to this clade. The phylogeny revealed that lions split from their sister species leopard ~3 Mya and the divergence time between snow leopards and the clade including lions and leopards was estimated to be ~5 Mya. Our study revealed that the morphology-based subspecies designation for both lions and tigers is largely not valid. The estimated tMRCA of 2.9 Mya between Barbary lions and Sub-Saharan African lions depicts the restriction of female-mediated gene flow between the lion populations in the backdrop of the habitat fragmentation taking place from late Pliocene to early to mid-Pleistocene creating islands of forest refugia in central Africa.

## Introduction

The charismatic big cats: tiger (*Panthera tigris*), lion (*Panthera leo*), leopard (*Panthera pardus*), snow leopard (*Panthera uncial*), and jaguar (*Panthera onca*) are grouped into the subfamily Pantherinae (Bagatharia, *et al.,* 2013). All of these big cats are endangered and are found in small fragmented populations in the world. Tigers are given an endangered status by IUCN (IUCN, 2015). They are currently found in Siberian taiga and grasslands (Siberian tiger, *P. t. altaica*), small pockets in Southeast Asia (Indochinese tiger, *P. t. corbetti,* and Malayan tiger, *P. t. jacksoni*), Southern China (South China tiger, *P. t. amoyensis*), island of Sumatra (Sumatran tiger, *P. t. sumatrae*), and the mangrove swamps of Indian subcontinent (Bengal tiger, *P. t. tigris*) (Luo, *et al.’* 2004). Lions are given a vulnerable conservation status (IUCN, 2015). They are found in two widely isolated geographical areas: various parts of Africa (African lion), and the Gir forest of Southwestern Gujarat, India (Asiatic lion) (Hemmer, 1966). While all Asiatic lion belong to the same subspecies (*P. l. persica*), the African lions are grouped into five separate subspecies: Barbary lion (Northern Africa, *P. l. leo*), Senegal lion (West-central Africa, *P. l. senegalensis*), Masai lion (East African lion, *P. l. nubica*), Katanga lion (Southwest Africa, *P. l. bleyenberghi*), and Transvaal lion (Southeast Africa, *P. l. krugeri*) (Hemmer, 1966). Leopards are designated as near threatened by IUCN (IUCN, 2015). These charismatic cats have a wide range of distribution from in sub-Saharan Africa to Southeast Asia and Siberia. There are currently nine recognized subspecies of leopard: African leopard (*P. p. pardus*), Arabian leopard (*P. p. nimr*), Persian leopard (*P. p. saxicolor*), Indian leopard (*P. p. fusca*), Sri Lankan leopard (*P. p. kotiya*), Indochinese leopard (*P. p. delacouri*), North-Chinese leopard (*P. p. japonensis*), Siberian leopard (*P. p. orientalis*), and Javan leopard (*P. p. melas*) (Uphyrkina, *et al.,* 2001). Snow leopards are listed as endangered in IUCN Red List of Threatened Species (IUCN, 2015). They are native to mountains of Central and South Asia. Jaguars are nearly endangered according to IUCN Red List (IUCN, 2015) and they are native to southern North America, Central and South America. There are three recognized subspecies of jaguar: Amazon jaguar (*P. o. onca*), Mexican jaguar (*P. o. hernandesii*), and Brazilian jaguar (*P. o. palustris*) (Seymour, 1989).

Reconstruction of the phylogenetic history of these charismatic cats can provide us a plethora of information about their species, subspecies, and population genetic status, which can be helpful for the conservation of these threatened animals.

Several morphological, biochemical, and molecular approaches have been employed to resolve the phylogenetic history of the Pantherine cats such as morphological (Salles, 1992), cytogenetic (Wurster-Hill and Gray, 1975), immunological (Collier and O'Brien, 1985), biochemical (Slattery, *et al.,* 1994), sex-chromosome based (Slattery and O'Brien, 1998), chemical (Bininda ‐ Emonds, Decker ‐ Flum and Gittleman, 2001), and molecular genetic (Bagatharia, *et al.,* 2013; Johnson, *et al.,* 2006; Johnson and O’brien, 1997; Wei, *et al.,* 2011). In spite of several trials to reconstruct the phylogenetic history of Pantherine cats, their phylogeny has remained largely unresolved. Johnson et al. (2006) grouped snow leopards with tigers based on 18’853 bp of nuclear DNA concatenated data (Johnson, *et al.,* 2006). In contrast, a later phylogenetic analysis grouped snow leopards with lions based on their complete mtDNA sequences (Wei, *et al.,* 2011). They estimated that *N. nebulosa* shared a common ancestor with other Pantherine cats about 8.66 Mya and that the leopards shared a common ancestor with the lion-snow leopard clade about 4.35 Mya. Recently, another mitogenomic study grouped lion and leopards together and placed snow leopard as the sister taxa to this clade (Bagatharia, *et al.*, 2013). Although *Panthera* phylogeny has been reconstructed multiple times, none of these studies used multiple subspecies of Pantherine cats. So, the subspecies anomaly remained unresolved.

All previous phylogenetic studies were performed using 5‐10 Pantherine cats, taking one individual from each species, which made it difficult to assess the phylogenetic tree at a high resolution and determine the subspecies level genetic diversity. In the present study we have used complete mitochondrial DNA sequences excluding the control region (15,489bp) from 43 publicly available Pantherine taxa including multiple individuals and subspecies of big cats to resolve the phylogenetic history of the gennus *Panthera* and to genetically validate the ‘subspecies’ status of various Pantherine taxa. This study thus offers an unparalleled and in-depth view of Pantherine phylogeny and genetically assess the ‘subspecies’ question that has remained elusive till date, which can aid in providing more effective conservation measures for these charismatic big cats.

## Methods

### Molecular phylogenetic analysis

We retrieved 43 publicly available Pantherine mitogenomes including all available subspecies of six charismatic cats: tiger, lion, leopard, snow leopard, jaguar, and Neofelis from GenBank (Benson, *et al.*, 2004) (Table 1). The mitogenome sequences were first aligned using Clustal Omega online server (Sievers, *et al.*, 2011). The fasta alignment of the complete mitogenome sequences was then exported to MEGA v6.06, where the control region (D-loop) sequences were eliminated. Pair-wise genetic distances among the Pantherine taxa were calculated in MEGA v6.06 (Tamura, et al. 2013). A model test was performed to determine the best fitting nucleotide substitution model for the dataset, using Akaike’s information criteria with correction (AICc) and Bayesian information criteria (BIC) using jModelTest v2.1.4 (Guindon and Gascuel, 2003). The alignment of 43 Pantherine taxa was used for maximum likelihood phylogenetic reconstruction, with 1000 bootstrap replicates, using MEGA v6.06. Bayesian phylogenetic tree was reconstructed using MrBayes v3.2.5 (Huelsenbeck and Ronquist, 2001).

**Table 1.** Gen Bank IDs, common, and scientific names of the Pantherine samples used in this study.

| Species | Common name | Subspecies | Common name | Gen Bank IDs |
| --- | --- | --- | --- | --- |
| 1. *Panthera tigris* (N = 13) | Tiger | a) *Panthera tigris amoyensis* (N= 2) | Amoy tiger | HM589214, HM589215 |
|  |  | b) *Panthera tigris altaica* (N= 4) | Siberian tiger | HM185182, JF357973, JF357974, KF297576 |
|  |  | c) *Panthera tigris tigris* (N= 3) | Bengal tiger | JF357967, JF357968, NC_010642 |
|  |  | d) *Panthera tigris sumatrae* (N= 1) | Sumatran tiger | JF357969 |
|  |  | e) *Panthera tigris jacksoni* (N= 1) | Malayan tiger | KJ508413 |
|  |  | f) *Panthera tigris corbetti* (N= 1) | Indochinese tiger | KJ508412 |
|  |  | g) Not mentioned (N= 1) | Not mentioned | KP202268 |
| 2. *Panthera leo* (N = 16) | Lion | a) *Panthera leo leo* (N= 2) | Barbary lion | KF776494, KF907306 |
|  |  | b) *Panthera leo persica* (N= 4) | Asiatic lion | JQ904290, KC834784, KP001501, NC_018053 |
|  |  | c) *Panthera leo senegalensis* (N= 2) | West African lion | KP001502, KP001497 |
|  |  | d) *Panthera leo azandica* (N= 1) | Congo lion | KP001506 |
|  |  | e) *Panthera leo nubica* (N= 3) | Masai lion | KP001495, KP001498, KP001499 |
|  |  | f) *Panthera leo bleyenberghi* (N= 2) | Southwest African lion | KP001504, KP001505 |
|  |  | g) *Panthera leo krugeri* (N= 1) | Transvaal lion | KP001500 |
|  |  | h) Not mentioned (N= 1) | Not mentioned | NC_028302 |
| 3. *Panthera pardus* (N = 5) | Leopard | a) *Panthera pardus pardus* (N= 1) | African leopard | KP001507 |
|  |  | b) *Panthera pardus japonensis* (N= 2) | North-Chinese leopard | EF551002, KJ866876 |
|  |  | h) Not mentioned (N= 2) | Not mentioned | KP202265, NC_010641 |
| 4. *Panthera uncia* (N = 3) | Snow leopard | NA | NA | EF551004, KP202269, NC_010638 |
| 5. *Panthera onca* (N = 4) | Jaguar | NA | NA | KF483864, KM236783, KP202264, NC_022842 |
| 6. *Neofelis nebulosa* (N = 2) | Clouded leopard | NA | NA | KP202291, NC_008450 |

### Dating the phylogenetic tree

For dating, we used a Bayesian Markov Chain Monte Carlo approach (MCMC) implemented in BEAST v1.8.2 (Drummond, *et al.*, 2012). The BEAST input file was generated using BEAUTi v1.8.2 (Drummond, *et al.*, 2012). An uncorrelated lognormal relaxed molecular clock was used to allow the evolutionary rate to vary from branch to branch. Previous studies have shown that has showed that relaxed molecular clock provides better estimate of time to most recent common ancestor (tMRCA) over the strict molecular clock that does not allow the evolutionary rate to vary among branches (Drummond, *et al.*, 2006). SRD06 model of nucleotide substitution was used to partition the nucleotide data by codon position. The rate of evolution was calibrated using lognormally distributed priors with lognormal means of zero and lognormal standard deviation of 0.56. Since lognormally distributed priors sample values from the more distant past more frequently than recent values, it performs better than both normally distributed and exponentially distributed priors, when using fossil calibration points for dating a phylogenetic tree (Ho, 2007). We offset the lognormal distributions of the priors at the internal node of *Panthera-Neofelis* split by 7 Mya, so that the median values of the sampled distributions were equal to and the mean values were slightly greater than the split date of ~8 Mya between the two. This calibration method has been previously employed for reconstructing phylogenetic history of other mammalian subfamily level taxa (Bjork, *et al.*, 2011; Das, *et al.*, 2014; Raaum, *et al.*, 2005). Yule speciation process, which has been shown to be more appropriate when analyzing sequences from different species, was used for the divergence estimation (Stone, *et al.*, 2010).

MCMC simulation ran for 10 million generations, sampling every 500 steps. 10% of the trees were removed as ‘burnin’. The maximum clade credibility (MCC) tree was identified and annotated using TreeAnnotator v1.8.2 (Drummond, *et al.*, 2012). Nodes with posterior probabilities exceeding 90% (P > 0.9) were used for the tree building. The MCC tree generated by TreeAnnotator v1.8.2 was visualized and graphically represented using FigTree v1.4.2 (http://tree.bio.ed.ac.uk/software/figtree). The posterior estimates of the parameters sampled by the Markov Chain was summarized in Tracer v1.6 (Rambaut and Drummond, 2013). tMRCA means, medians, and 95% highest posterior density (95% HPD) intervals (all in Myr) were also calculated in Tracer v1.6.

## Results

### Sequence divergence and the subspecies dilemma

Among lions, identical nucleotide sequences were observed among *P. l. senegalensis* (KP001502), *P. l. azandica* (KP001506), *P. l. nubica* (KP001495), and *P. leo* (NC_028302), raising questions about the validity of the different subspecies status for these individuals. The two Barbary lions (KF776494 and KF907306) are genetically most distant from the rest of the clade, showing 2.2 - 3% divergence of the mitogenome. The remaining lion subspecies shows 0 - 0.6% genetic divergence among each other. Our results thus indicate that the morphological variation among the different African lion populations may not be enough to consider them as different ‘subspecies’.

The validity of the subspecies status is also questionable in case of the tigers with *P. t. amoyensis* (HM589215) and *P. t. altaica* (HM185182) showing identical nucleotide sequences. The mean tiger mitogenome divergence was 0.5% and the largest divergence was observed between the Amoy tigers (HM589214 and HM589215) and the rest of the tree (1.2%). Thus, similar to lions, most of the tiger subspecies are also potentially not valid genetically.

Leopard subspecies show 0 – 0.9% genetic divergence among each other with the highest divergence between *P. p. pardus* (KP001507) and *P. p. japonensis* (EF551002) (0.9%).

### Phylogenetic analyses

The maximum likelihood (ML) tree was constructed using Hasegawa-Kishino-Yano model of nucleotide substitution with gamma distribution and invariable sites (HKY+G+I), which was selected to be the best fitting model for the dataset by jModelTest v2.1.4. The ML based phylogeny grouped lions (N = 16) and leopards (N = 5) together with 100% bootstrap support and placed show leopards (N = 3) as the sister taxa to this clade (100% bootstrap support) (Figure 1). Jaguars (N = 4) were designated to be the sister taxa of the lion-leopard-snow leopard clade (Figure 1). The deepest root within the lions was observed between the Barbary lions (*P. l. leo*) and the rest of the subspecies with 100% bootstrap support (Figure 1). For the tigers (N = 13), the deepest root was observed between the Amoy tigers and the rest (100% bootstrap support) (Figure 1). Similar grouping of the Patherine taxa was revealed in the Bayesian phylogenetic tree (Figure 2). Overall, the phylogenetic analyses revealed a dendrogram: *Neofelis nebulosa* (*Panthera tigris* (*Panthera onca* (*Panthera uncia* (*Panthera leo*, *Panthera pardus*)))).

**Figure 1.**
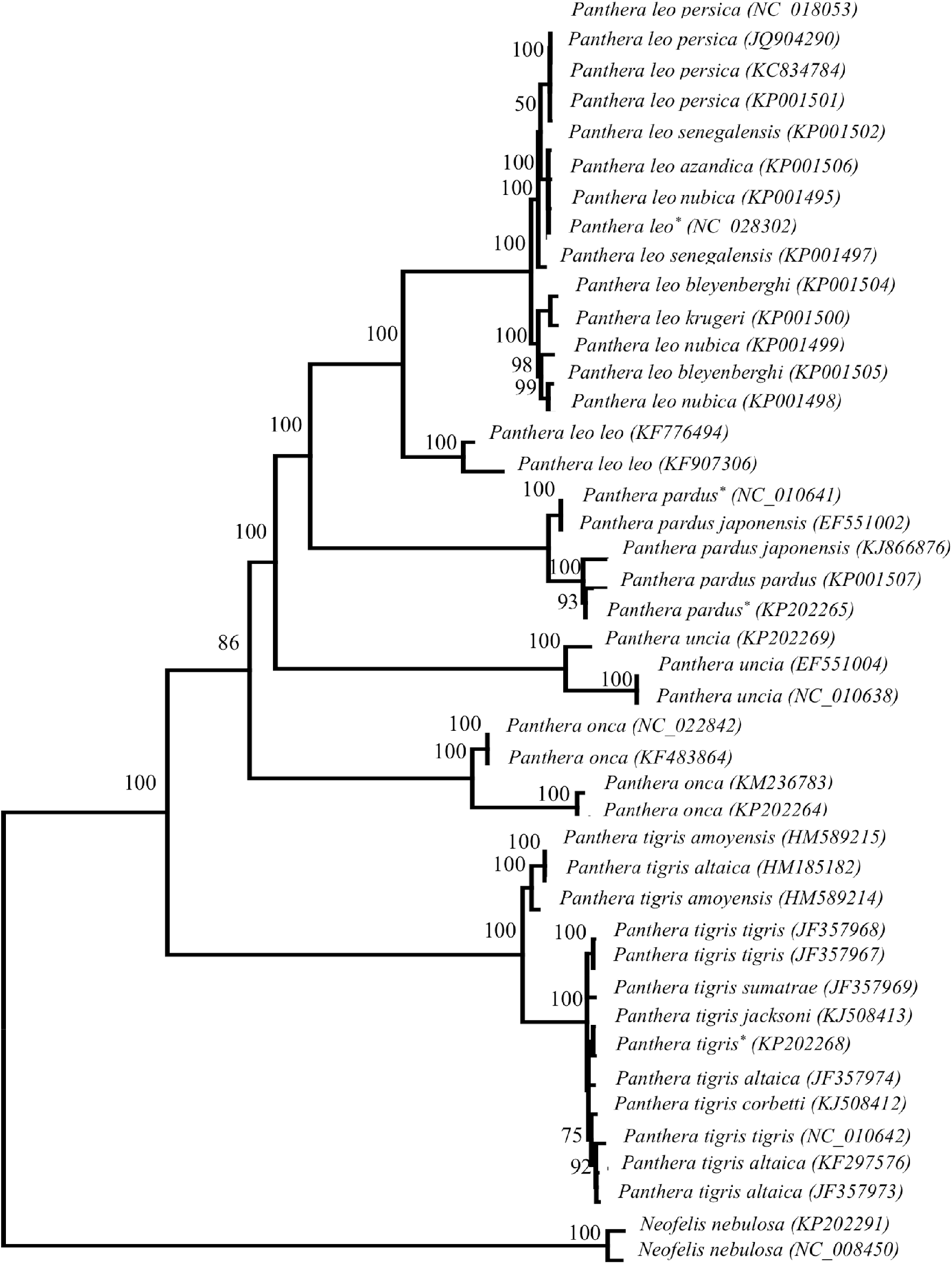
Consensus maximum likelihood tree (HKY+I+G model) of 43 Pantherine taxa. Percentage of bootstrapped replicates supporting each node (out of 1,000 bootstraps) are shown. The nodes with less than 50% bootstrap support are not shown. The *Neofelis* cats were used as the outgroup to the genus *Panthera.* The individuals with unknown subspecies status are marked with an asterisk.

**Figure 2.**
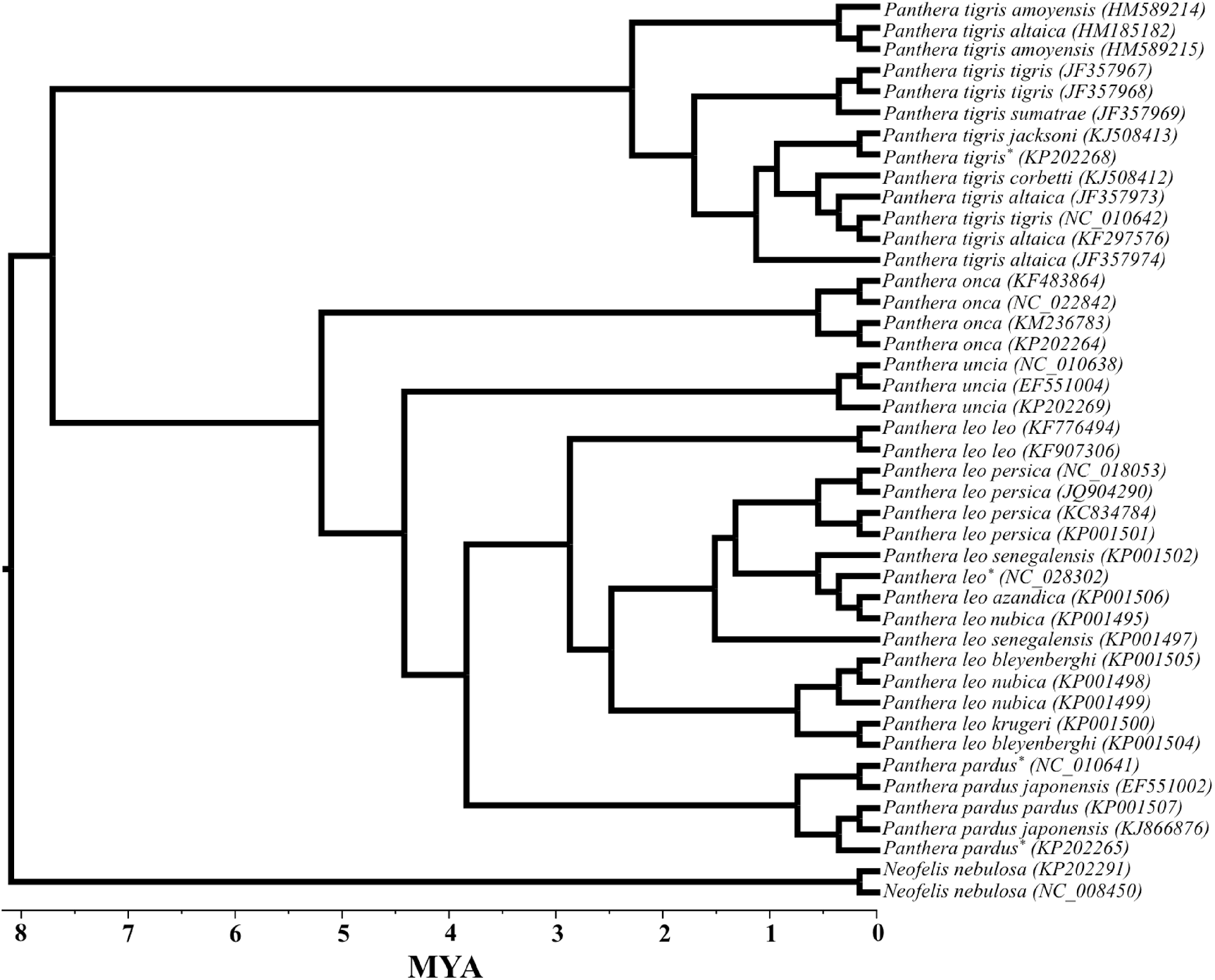
Phylogeny of 43 Pantherine taxa based on complete mtDNA sequences excluding the control region. The dates were inferred using a Bayesian MCMC approach implemented in BEAST with *Neofelis-Panthera* divergence offset by 7 Myr to approximate a 8 Mya date for *Neofelis-Panthera* split. Individuals with uknown subspecies status are indicated with an asterisk.

### Estimates of divergence dates

As mentioned before (see methods) a single calibration point of *Neofelis-Panthera* split of 8 Mya was used to estimate the divergence time. Similar to the ML tree, the Bayesian tree also grouped lions and leopards together and placed snow leopard as the sister taxa to this clade. The MCMC analysis employed in BEAST revealed that lions split from its sister species leopard 3.45 Mya (1.75 – 5.17) (Table 2). The divergence time between snow leopards and the clade including lions and leopards was estimated to be 4.97 Mya (2.9 -7.24) (Table 2). Tigers appeared to split from the rest four charismatic big cats 7.47 Mya (4.97 – 9.24) (Table 2). The deepest roots within the tigers was estimated to be ~2 Mya between the Amur tiger and the remainder of the tiger clade (Table 2). As revealed by the ML tree, the deepest split within lions was between the Barbary lions and the remainder of the subspecies (including all Sub-Saharan African and Asian subspecies) (2.87 Mya). The divergence between the Asian lions from the East-Central African subspecies is relatively new 1.39 Mya (0.63 – 4.2) (Table 2).

**Table 2.** tMRCA dates with confidence intervals.

| **Taxon divergence** | **tMRCA mean<sup>a</sup>** | **95% HPD<sup>b</sup>** | **Median** |
| --- | --- | --- | --- |
| *Neofelis* - *Panthera* | 8.03 | 7.222 - 9.213 | 7.894 |
| Tiger - Lion/Leopard/Snow leopard/Jaguar | 7.467 | 4.972 - 9.236 | 7.655 |
| Jaguar - Lion/Leopard/Snow leopard | 6.708 | 3.923 - 8.850 | 6.94 |
| Snow leopard - Lion/Leopard | 4.968 | 2.9 - 7.243 | 4.852 |
| Lion - Leopard | 3.45 | 1.746 - 5.166 | 3.398 |
| Deepest root within tiger | 2.333 | 0.932 - 3.868 | 2.252 |
| Asian and East-Central African lions | 1.39 | 0.63 - 4.2 | 0.85 |

<sup>a</sup>Time to most recent common ancestor, in Myr.<sup>b</sup>95% highest posterior density

## Discussion

### mtDNA based phylogeny: advantages of large sample size

For the last two decades mtDNA is being used to reconstruct Pantherine phylogeny. However, all previous phylogenetic analyses in this regard were employed using 5 – 10 Pantherine taxa. Moreover, none of them included multiple subspecies and/or individuals from a species. Consequently, the deepest root within the species remained largely unknown. Also, due to low resolution, the Pantherine species were often wrongly grouped, often generating contrasting results. For instance, Johnson et al (2006) grouped snow leopards with tigers and Wei et al. (2011) grouped them with lions (Johnson, *et al.*, 2006; Wei, *et al.*, 2011).

We used 43 Pantherine taxa in this study, including individuals and subspecies from all charismatic big cats. Complete mtDNA sequences excluding the control regions (15,489bp) were used for phylogenetic analysis. The control regions were excluded in this study on two major accounts: i) this area of the mitogenome is very difficult to align across the big cats and ii) the repeat sequences found in this region results in different mitogenome sizes across the Pantherine cats (Bagatharia, *et al.*, 2013). Therefore several recent phylogenetic studies on Pantherine cats have either included only the protein coding genes or the complete mtDNA sequence excluding the control region (Bagatharia, *et al.*, 2013; Wei, *et al.*, 2011). Proper and systematic sampling method employed here aids to the robustness of the analyses by increasing the statistical power. Both ML and Bayesian based phylogeny revealed the same grouping of all Pantherine taxa.

Overall, the current study revealed a dendrogram: *Neofelis nebulosa* (*Panthera tigris* (*Panthera onca* (*Panthera uncia* (*Panthera leo*, *Panthera pardus*)))). The inclusion of multiple individuals and subspecies from all species and incorporation of jaguars has helped to reconstruct hitherto unresolved Pantherine phylogeny with high confident. Snow leopards are genetically distant from the tigers with 7.5 – 8.1% sequence divergence and therefore grouping them with tigers is potentially an artifact of improper and inadequate sampling (Johnson, *et al.*, 2006). Lions are genetically close to both leopards (4 – 5% sequence divergence) and snow leopards (5.5 – 6.1% sequence divergence). So, if multiple individuals from all species are not included in the analyses, lions can be, at random, wrongly grouped with snow leopards, and this is what has potentially occurred in case of the phylogeny described in Wei et al. (2011) who used only six Pantherine taxa in their study (Wei, *et al.*, 2011). Unlike the aforementioned studies, the recent-most Pantherine tree described in Bagatharia et al. (2013) revealed correct species level phylogeny by grouping lions and leopards together and placing snow leopards as the sister species to the lion-leopard clade. But the low sample size (six Pantherine cats) of the study and the exclusion of jaguars in all analyses resulted in a low-resolution phylogeny.

### Subspecies anomaly

Our study raised question on ‘subspecies’ designation based on morphology. In most cases we found that the current ‘subspecies’ designation is not genetically valid and needs serious taxonomic change.

For instance, current lion subspecies status is largely based on morphological characteristics mentioned in Hemmer (1966) (Hemmer, 1966). The current study raises questions about the authenticity of the subspecies status of various Sub-Saharan African lion subspecies based on morphology. Although currently geographically isolated, *P. l. senegalensis* (KP001502), *P. l. azandica* (KP001506), *P. l. nubica* (KP001495), and *P. leo* (NC_028302) share identical nucleotide sequences. In contrast, the two *P. l. senegalensis* individuals (KP001497 and KP001502) show 0.3% nucleotide divergence. Similar nucleotide divergence (0.4%) was observed between the two *P. l. bleyenberghi* individuals (KP001504 and KP001505). These results indicate recent geographical isolation of Sub-Saharan African lion populations, who potentially have not yet accumulated subspecies level genetic divergence. For decades’ authors have debated on the exact number of lion subspecies (eight vs. ten). Our study conclusively revealed that there are potentially only three ‘subspecies’ of lion currently exists in this planet: North African Barbary Lion, Sub-Saharan African Lion, and Asiatic Lion. The genetic variations observed among the Sub-Saharan African Lions are potentially mere individual variations.

The validity of the subspecies status is also questionable in case of the tigers with *P. t. amoyensis* (HM589215) and *P. t. altaica* (HM185182) showing identical nucleotide sequences and two *P. t. altaica* individuals (HM185182 and JF357973) showing high (1.1%) nucleotide divergence. The aforementioned results clearly indicate more individual level variation (intra-population) than subspecies level variation (interpopulation), raising questions about the validity of morphology-based subspecies designation. It can be speculated that if more samples were analyzed, individuals from different populations would have grouped together disobeying their current ‘subspecies’ status.

### Migration and isolation of Pantherine taxa

Tigers potentially split from other five big cats sometime in late Miocene. As speculated in previous studies, *Panthera* species such as lions potentially migrated into America and Africa from Asia in the Pliocene (Werdelin and Lewis, 2005). The migration path of the Pantherine cats mentioned in Johnson et al. (2006) fits with the divergence times of Pantherine cats mentioned in this study (Johnson, *et al.*, 2006). Snow leopards potentially got isolated from lions and leopards in early Pliocene (~5Mya), when these cats started migrating to Africa. The estimated tMRCA of 2.9 Mya between Barbary lions and the Sub-Saharan African lions is an indication of potential restriction of female-mediated gene flow between these populations. The habitat fragmentation from late Pliocene to early to mid-Pleistocene resulted in the islands of forest refugia in central Africa, isolating Barbary lions from other lion populations. However, migration and intermixing potentially continued among the Sub-Saharan African subspecies till forests were converted into the savannas during late Pleistocene. Additionally it is interesting to note that be noted here that the mitogenome divergence among the lion populations potentially began at the same time as other African mammals such as chimpanzees, bonobos, and gorillas. Overall, this study provides a holistic and in-depth view of Pantherine phylogeny, indicating the importance of using large number of samples for confident resolution of mammalian phylogenetic history.

